# drFrankenstein: An Automated Pipeline for the Parameterisation of Non-Canonical Amino Acids

**DOI:** 10.64898/2026.03.16.712088

**Authors:** Eugene Shrimpton-Phoenix, Evangelia Notari, Christopher W. Wood

## Abstract

The incorporation of non-canonical amino acids (ncAAs) is a powerful strategy for introducing novel chemical functions into proteins. Molecular dynamics (MD) simulations are essential for understanding the structural and dynamic effects of these modifications, yet the creation of accurate force field parameters for ncAAs remains a significant bottleneck. Current parameterisation methods are often inaccurate or computationally expensive.

To address this, we present drFrankenstein, an automated pipeline for generating AMBER force field parameters for ncAAs. drFrankenstein is a robust and accessible tool that streamlines the parameterisation workflow, enabling the routine use of MD simulations to study the behaviour of ncAA-containing proteins.

## Introduction

Protein engineering has been significantly advanced by genetic code expansion technology, which enables the incorporation of non-canonical amino acids (ncAAs) into protein structures (Shandell *et al*., 2021). This capability has unlocked a diverse range of applications, such as increasing the stability of peptide-based drugs (Miranda *et al*., 2008) and the prevention immunogenic responses (Schultz *et al*., 2018). Furthermore, the incorporation of ncAAs into enzyme active sites provides access to novel chemical reactivities not seen in nature (Burke *et al*., 2019; Trimble *et al*., 2022).

Recently, the field of protein design has seen a seismic shift as the result of deep-learning methods. Previously intractable problems such as protein-folding have become routine (Jumper *et al*., 2021; Watson *et al*., 2023; Hayes *et al*.). Unfortunately, structural data for ncAA containing proteins is scarce. This limits the effectiveness of deep-learning based techniques for designing proteins that contain ncAAs. Instead, researchers must rely on physics-based methods such as molecular dynamics (MD) simulations to probe the structure, function and interactions of ncAA containing proteins.

To perform a MD simulation on a system, each atom, residue and molecule must have force field parameters associated with it. These parameters describe how bonds, angles, dihedrals and non-bonded interactions behave throughout the simulation. While robust force field parameters exist for the 20 canonical amino acids (Brooks *et al*., 1983, 2009; Abraham *et al*., 2015; Case *et al*., 2023), no such set exists for ncAAs. As a result, before a researcher performs MD simulations on a ncAA-containing protein, they must create some supplemental parameters that describe their ncAA of interest.

At the most basic level, these parameters can be created “by-analogy” (Vanommeslaeghe *et al*., 2010; Zoete *et al*., 2011). This approach simply assigns parameters to new molecules by finding appropriately close parameters in pre-existing forcefields. While this approach is extremely fast, defining parameters by analogy could be inaccurate. For exotic ncAAs with functional groups that do not exist in the pre-existing forcefield, the resulting parameters will provide a poor description of its dynamics (Wildman *et al*., 2016).

Alternatively, quantum mechanical (QM) methods are employed to derive parameters for new molecules from scratch (Mayne *et al*., 2013). This *ab initio* approach has the potential to create parameters that better describe the dynamics of an exotic species than “by-analogy” methods. The considerable downside of these methods is that they can be extremely computationally expensive and often require non-standardised manual steps.

To address these challenges, we present drFrankenstein (available at https://github.com/wells-wood-research/drFrankenstein), a comprehensive and fully-automated pipeline for parameterising ncAAs for the AMBER force field. It handles the entire workflow; from terminal capping and conformer generation to torsion scanning, charge calculations and parameter fitting (Figure 1). To simplify setup and ensure reproducibility, the entire calculation is controlled via a single YAML input file.

**Figure 1:**
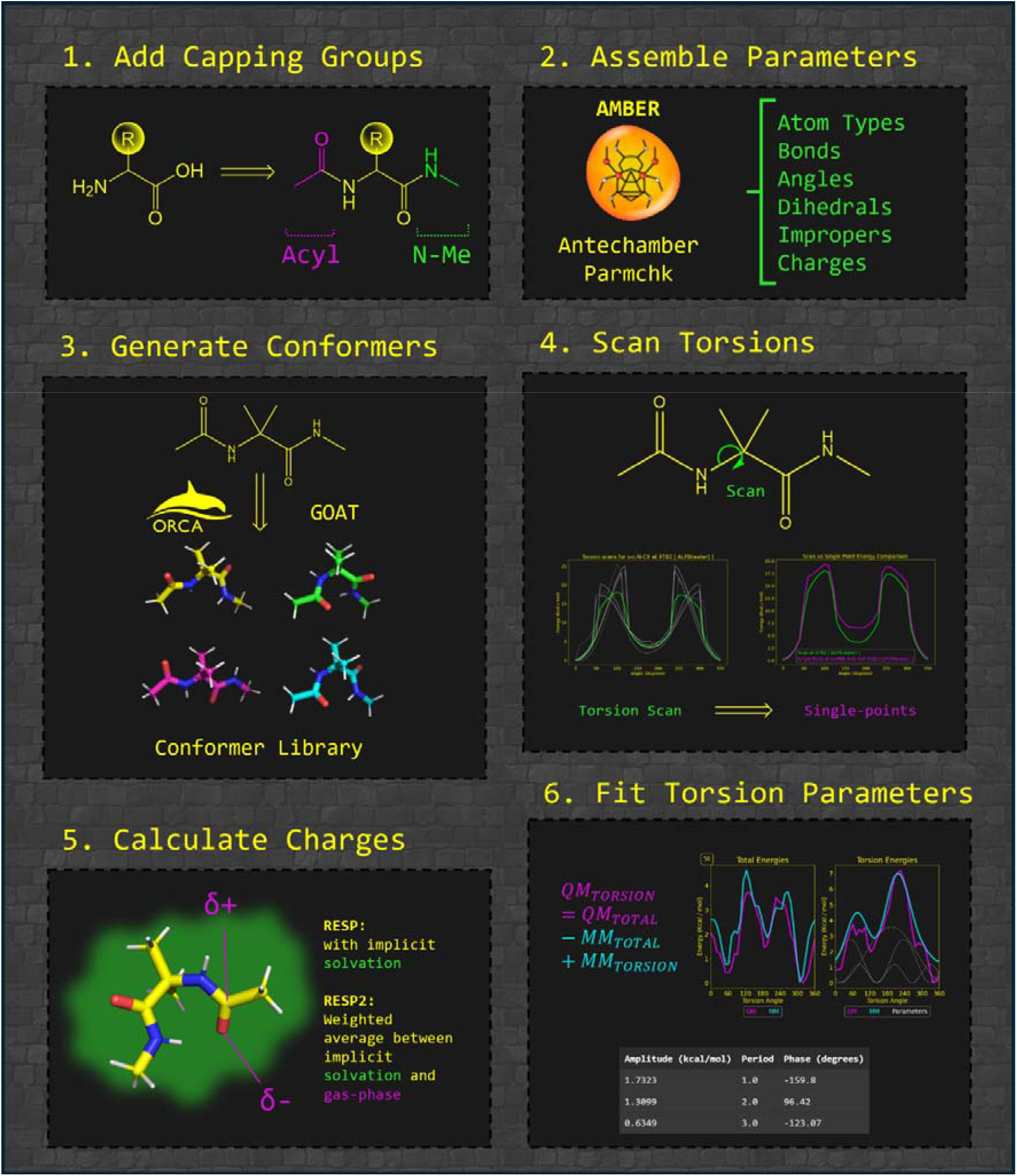
The main protocols performed by drFrankenstein: First capping groups are added to C and N termini. Then a set of starting parameters are created. Next, a set of conformers are generated. These conformations are then used as inputs for torsion scanning and charge calculation steps. Finally, MD parameters are fit to the torsion scans resulting in a final parameter set.

## Results

### Workflow

Initially, drFrankenstein adds acetyl capping groups to any user-defined N-termini and N-Methyl capping groups to any user-defined C-termini. This ensures that parameters created describe the interaction between the ncAA of interest and adjacent backbone atoms of the protein chain.

Next, an initial set of “by-analogy” parameters are generated using Antechamber (Case *et al*., 2023). The bond, angle, improper and non-bonded parameters in these “by-analogy” parameters will be used in drFrankenstein’s final parameters. In subsequent steps, the torsion and charge parameters will be recalculated using QM-based methods.

Once capping groups have been added, a library of low energy conformers of the ncAA are generated using ORCA’s global local minima searching tool GOAT (Neese, 2012). These conformers will be used as input geometries for the subsequent torsion scanning and charge calculation steps.

The next step is to perform torsion scans; drFrankenstein automatically detects all rotatable bonds in the ncAA of interest. For each rotatable bond, grouped by atom type, drFrankenstein uses ORCA (Neese, 2012) to perform a pair of relaxed torsion scans in the forwards and backwards direction using an interval of 10°. Scans are performed using different conformers as starting points, this allows for different energy pathways to be explored. The final energy profile is constructed as the geometric average of each energy profile generated. Optionally, the energy profile of each torsion scan can be re-evaluated using a series of single-point energy calculations along the scan trajectory. By using a relatively inexpensive QM method to perform the torsion scan and a more accurate method to perform the single-points, the user can maximise both the accuracy and speed of this step.

The restrained electrostatic potential (RESP) protocol (Woods and Chappelle, 2000) has been implemented in drFrankenstein: First a geometry optimisation is performed, followed by a single-point energy calculation on each input conformer using ORCA. The resulting wavefunction files are then passed to the program MultiWFN (Lu and Chen, 2012) to perform RESP charge fitting, which produces the final partial charges. This procedure is performed using several conformers of the ncAA, with the final partial charges calculated using a Boltzmann-weighted average. We have also implemented the RESP2 procedure (Schauperl *et al*., 2020), which performs the above charge calculation procedure with and without implicit solvent. The final partial charges are then calculated as a weighted average (60:40 solvated: gas-phase).

Once torsion scans and charge fitting procedures have been completed, the next step is to assemble a set of torsion parameters that accurately describe the energy profiles generated in the torsion scanning step. To parameterise an individual torsion, the energy profile is calculated by:

***Equation 1:*** *The calculation of QM*_*TORSION*_ *from* torsion scan *energy*.

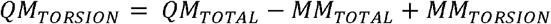

Where QM_TORSION_ is the energy associated with solely the torsion in question, and QM_TOTAL_ is the energy profile calculated in the torsion scanning step. MM_TOTAL_ is calculated by performing single-point energy calculations at the MM level, using the current iteration of the MM parameters, upon the trajectories generated by the torsion scanning step, which is performed using OpenMM (Eastman *et al*., 2024). MM_TORSION_ is extracted from the current iteration of the MM parameters directly as a linear combination of dihedral terms associated with the torsion in question.

Once QM_TORSION_ has been calculated, cosine functions are fitted to this energy profile using the Inverse Fast Fourier Transform. The amplitudes, periodicities and phase-shifts that describe the most significant of these cosine functions are then selected as parameters to describe the torsion.

The above process is performed iteratively over each torsion of interest several times, updating the MM parameters after each step. In this manner, each time QM_TORSION_ is calculated, the values of MM_TOTAL_ and MM_TORSION_ are evaluated in the context of the previously parameterised torsions. The order in which the torsions are parameterised is shuffled each iteration of this procedure. drFrankenstein applies a L2-dampening to the amplitudes to maintain stability during this process. After multiple iterations of this parameter fitting procedure, a set of torsion parameters that accurately describe the energy profiles generated in the torsion scanning step will have been created (Figure 2).

**Figure 2.**
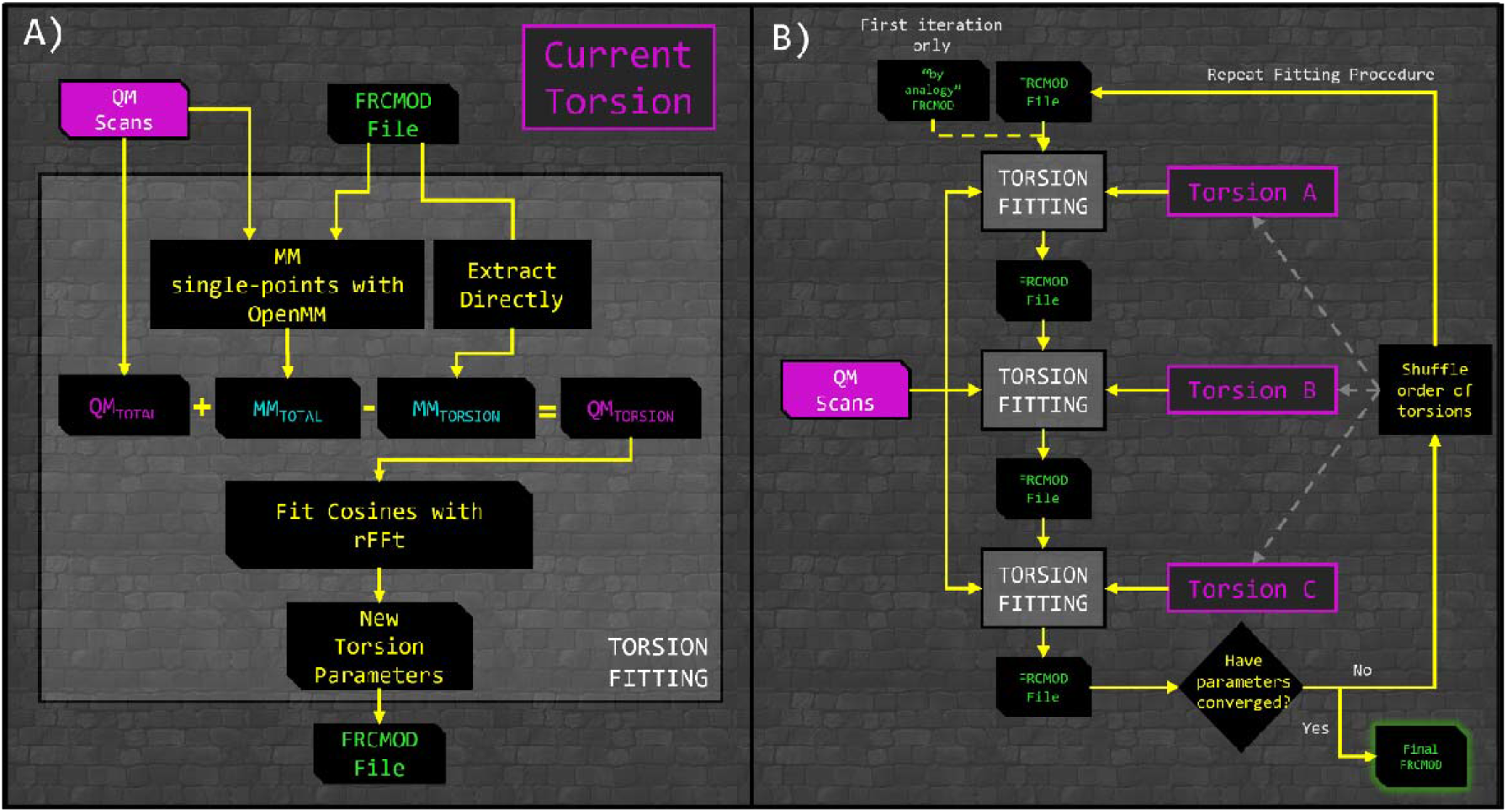
(A) Workflow diagram for the torsion parameter fitting process for a single torsion angle. (B) Workflow diagram for the iterative torsion parameter fitting process for all torsions: The per-torsion parameter fitting procedure shown in (A) is performed sequentially for each torsion angle. This sequence is performed iteratively, shuffling the order of the torsions until convergence criteria are met.

Once the parameter fitting process has completed, optional modifications to the parameter set are made, such as adding CMAP terms, and the duplication of torsion parameters to account for edge-cases where ncAAs are adjacent to each other or are terminal residues.

### Quality-of-life features

Great care has been put into the creation of drFrankenstein to ensure that it is reliable, interpretable and easy to use. To this aim, a single configuration file is passed to drFrankenstein, this both simplifies the setup of calculations and ensures reproducibility.

Once drFrankenstein has completed all of its calculations, it creates an interactive report that details the key results of its calculations at each step. This report can be used to check if the parameters generated are as the user expects. Additionally, each page of this report contains a plain-English explanation of the processes that drFrankenstein has performed, tailored to the user’s calculations. This explanation is accompanied by links to key citations associated with each step. The purpose of this is to aid in the writing of methods sections for academic publications.

drFrankenstein allows for any QM method to be used in each parameterisation step, provided it is in ORCA’s *(Neese, 2012)* extensive methods library. This allows users to use computationally cheaper QM methods, sacrificing some accuracy to dramatically decrease the computational cost or use sacrifice speed in order to ensure legacy compatibility.

### Use case 1: Creation of AMBER parameters for AIB

To demonstrate the use of drFrankenstein, we used it to create AMBER-compatible parameters for the ncAA 2-Aminoisobutyric acid (AIB). The incorporation of AIB into peptides has been shown experimentally to induce the formation of 3-10 helices (Kumar *et al*., 2022). The sterically constrained Phi and Psi angles of AIB are responsible for this behaviour, which we parameterised using drFrankenstein. Further details on the parameterisation of AIB can be found in **Supplementary Information**.

Using the parameters for AIB generated by drFrankenstein, we performed MD simulations using drMD (Shrimpton-Phoenix *et al*., 2025) on three short peptides, each containing three AIB residues (Figure 3A). These peptides have been experimentally shown to partially adopt 3-10 helix structures (Karle *et al*., 1994). As a control, we also ran the same simulations on the peptides where the AIB residues had been mutated to Alanine. To account for the relative instability of 3-10 helices, we ran production simulations at 200K.

**Figure 3.**
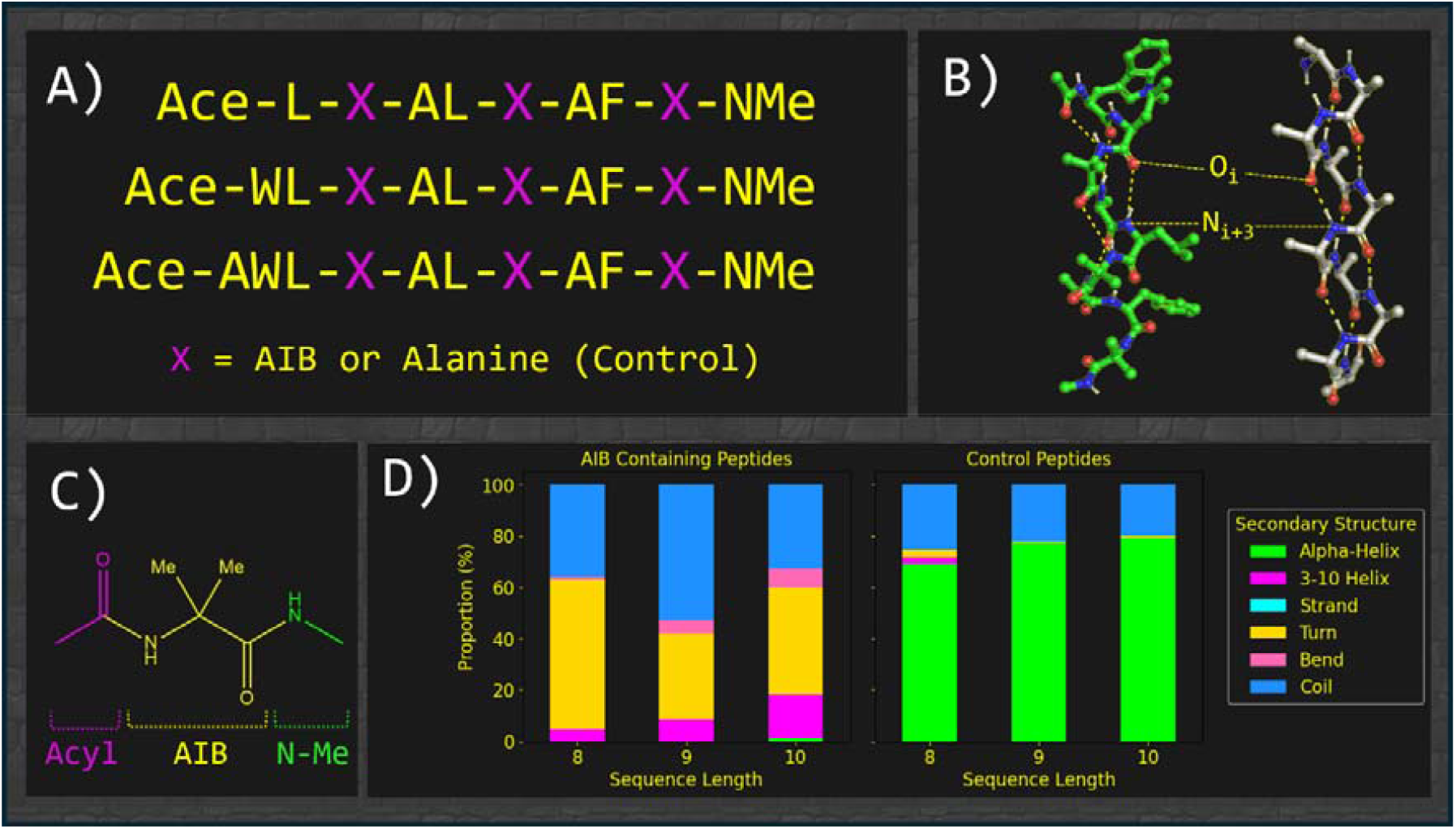
(A) Peptide sequences for our simulations. Each peptide was capped with Acyl and N-Methyl groups. (B) An idealised structure of a 3-10 helix (shown in white) and an MD snapshot of the AIB-containing octapeptide in which it adopts a partial 3-10 helix structure (shown in green). (C) The chemical structure of AIB, with capping groups. (D) Secondary structure proportions for both AIB-containing and control peptides, measured throughout 50 ns MD simulations.

Over the course of each 50 ns simulation, we observed partial 3-10 helix formation in the AIB-containing peptides. For the control peptides, we instead observed the formation of α-helices. This demonstrates that the drFrankenstein protocol is capable of generating forcefield parameters that reproduce experimentally observed behaviour.

### Use case 2: Creation of AMBER parameters for photo-caged tyrosine and GFP Chromophore

Green fluorescent protein (GFP) forms a complex with the anti-GFP “enhancer” nanobody (eNB). A key interaction between GFP and eNB is mediated by eNB Tyr37. In conjunction with some binding site redesign, the replacement of Tyr37 with the ncAA ortho-nitrobenzyl tyrosine (ONBY) has been shown to inhibit the formation of the GFP-eNB complex (O’Shea *et al*., 2022).

To simulate the interaction between GFP and eNB^Y37ONBY^, we are required to create two sets of ncAA parameters: One for the ONBY itself, and one for the chromophore CRO located at the core of GFP. Details on the parameterisation of ONBY and CRO can be found in **Supplementary Information**.

We performed short (50 ns) MD simulations on the eNB-GFP complex with and without the ONBY ncAA at position 37. In the simulation without ONBY, we observe a stable hydrogen bonding interaction between Tyr37 on eNB and Arg164 on GFP. In our simulation containing the ONBY residue, this hydrogen bonding interaction was disrupted by the displacement of Arg164 by the steric bulk of the ONBY protecting group (see Figure 4B & D).

**Figure 4.**
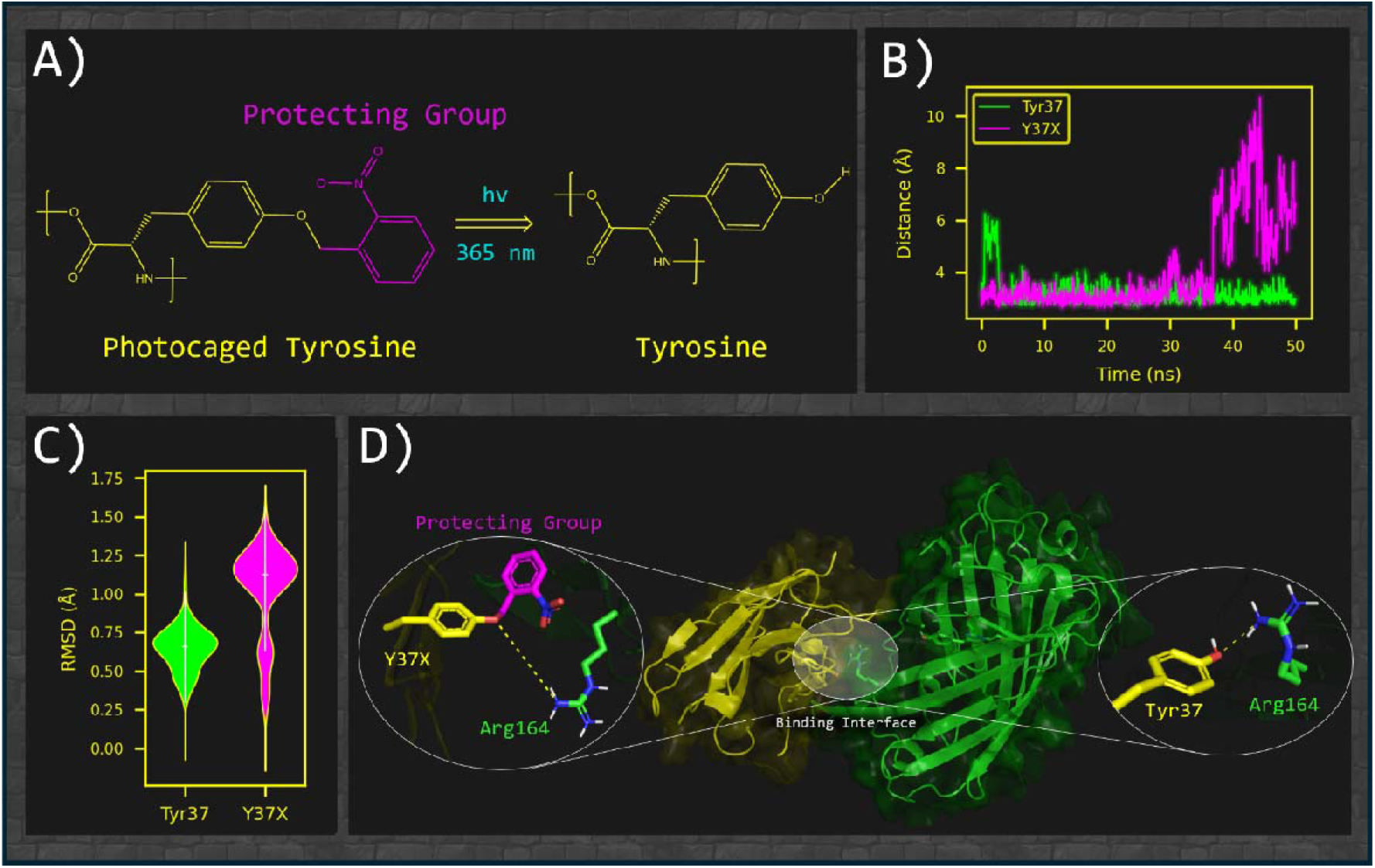
(A) The chemical structures of ncAA ONBY and its light-activated degradation to tyrosine. (B) The time-evolved interaction distance with GFP[Arg164-NH2] for eNB[Tyr37-OH] and eNB[Y37ONBY-OH]. (C) Violin plots of the RMSD of the ncAA CRO for our simulations. (D) Structure of the GFP (green) – eNB (yellow) complex. When unprotected, eNB[Tyr37] forms a stable hydrogen bond with GFP[Arg164]. When protected by ONBY, this interaction is disrupted.

## Conclusions

drFrankenstein builds upon our previous program Stapline (Notari *et al*., 2025), which focuses on the parameterisation of stapled amino acids. drFrankenstein focuses instead on the broader field of ncAAs and has been thoroughly reworked to ensure robustness, ease of use and computational efficiency.

drFrankenstein is a robust pipeline capable of generating parameters compatible with the AMBER forcefield. By leveraging the extensive ORCA inputs library, drFrankenstein allows the user to choose the level of theory at each step, so that the user can balance their need for accuracy with computational cost. We have demonstrated that the parameters produced using a combination of GFN-XTB2 and rev2PDE def2-SVP D3BJ levels of theory is sufficient to reproduce experimentally observed behaviours in proteins. This is of great significance as the levels of theory used are orders of magnitude faster than those traditionally used. This allows for facile parameterisation of larger and more chemically diverse ncAAs.

## Acknowledgements

Eugene Shrimpton-Phoenix and Christopher W. Wood are supported by a BBSRC sLOLA award (BB/X003027/1). Evangelia Notari is supported by a PhD studentship from the UKRI funded EastBio Doctoral Training Partnership programme.

