## Supplementary Information for "drFrankenstein: An Automated Pipeline for the Parameterisation of Non-Canonical Amino Acids"

### Details on the parameterisation of AIB

Torsion scans were performed at the XTB2 level with the ALPB implicit solvation method. Torsion scan energy profiles were then reevaluated at the revPBE def2-SVP D3BJ level with the CPCM implicit solvation method. Partial charges were calculated using the RESP2 protocol with geometries optimised at the XTB2 level and charge distributions calculated at the revPDB def2-SVP D3BJ level. Torsion parameters were fit using a maximum of three cosine functions per torsion parameter.

### Details on the parameterisation of ONBY and CRO

For the parameterisation of ONBY, we chose not to optimise the parameters associated with the Phi and Psi torsions, instead we used the standard parameters from the AMBER19ff forcefield. This was done to ensure that our ONBY parameters do not interfere with the secondary structures of the protein. For CRO, there is no standard Phi/Psi angles due to its non-traditional backbone structure. As such, all torsion angles involving heavy atoms were parameterised.

Torsion scans were performed at the XTB2 level with the ALPB implicit solvation method. Torsion scan energy profiles were then reevaluated at the revPBE def2-SVP D3BJ level with the CPCM implicit solvation method. Partial charges were calculated using the RESP2 protocol with geometries optimised at the XTB2 level and charge distributions calculated at the revPDB def2-SVP D3BJ level. Torsion parameters were fit using a maximum of three cosine functions per torsion parameter.

### Details on MD simulations performed for case study 1

Input structures for the AIB-containing peptides were generated using the software tleap, available as part of the AMBERTOOLS package (Case *et al.*, 2023).

MD simulations were performed in our own simulation pipeline drMD (Shrimpton-Phoenix *et al.*, 2024). For each peptide, the following simulation protocol was applied: First, an energy minimisation step was performed for 1000 steps or until the system’s energy converged. Next a short (400 ps) simulation under the canonical (NVT) ensemble was performed, increasing the temperature from 0 K to 300 K in 100 K intervals. Next a short (100 ps) simulation was performed under the isothermal-isobaric (NpT) ensemble at 300 K. In the two previously described simulation steps, a position restraint was applied to each protein atom. The purpose of this restraint is to allow the positions of solvating water molecules to equilibrate. Next, a very short (10 ps) simulation was performed, without position restrains and under the NpT ensemble. In this simulation step, a shorter (0.5 fs) simulation timestep was used. This step was performed to ensure the simulation remained stable after the removal of the position restraints used in the previous two steps. Finally, a longer (50 ns) simulation was performed at 200 K and under the NpT ensemble.

All simulations were performed using the OpenMM simulation toolkit (Eastman *et al.*, 2017). The following methods were used in all MD simulations: Velocities of the system were calculated using the Langevin middle integrator (Zhang *et al.*, 2019). Non-bonded interactions were modelled using the Particle-Mesh Ewald method with a non-bonded cutoff of 10 Å (Darden *et al.*, 1993). Constraints were applied to all bonds between heavy atoms and hydrogen atoms using the SHAKE algorithm (Kr�utler *et al.*, 2001). Bonds and angles of water molecules were restrained using the SETTLE algorithm (Miyamoto and Kollman, 1992). For NpT simulations, the Monte-Carlo barostat was used to enforce a constant pressure of 1 bar. We treated all protons as having a mass of 4.03036 amu, the extra mass being subtracted from adjacent heavy atoms (Feenstra *et al.*, 1999). This, combined with hydrogen bond constraints allowed all simulations to be performed using a timestep of 4 fs, which roughly doubled our simulation speed.

### Details on MD simulations performed for case study 2

Input structures for the ONBY-containing enB^ONBY^ were generated using the software tleap, available as part of the AMBERTOOLS package (Case *et al.*, 2023).

MD simulations were performed in our own simulation pipeline drMD (Shrimpton-Phoenix *et al.*, 2024). For each peptide, the following simulation protocol was applied: First, an energy minimisation step was performed for 1000 steps or until the system’s energy converged. Next a short (400 ps) simulation under the canonical (NVT) ensemble was performed, increasing the temperature from 0 K to 300 K in 100 K intervals. Next a short (100 ps) simulation was performed under the isothermal-isobaric (NpT) ensemble at 300 K. In the two previously described simulation steps, a position restraint was applied to each protein atom. The purpose of this restraint is to allow the positions of solvating water molecules to equilibrate. Next, a very short (10 ps) simulation was performed, without position restrains and under the NpT ensemble. In this simulation step, a shorter (0.5 fs) simulation timestep was used. This step was performed to ensure the simulation remained stable after the removal of the position restraints used in the previous two steps. Next a mid-length (5 ns) equilibration step was performed under the NpT ensemble at 300 K. Finally, a longer (50 ns) simulation was performed at 300 K and under the NpT ensemble.

All simulations were performed using the OpenMM simulation toolkit (Eastman *et al.*, 2017). The following methods were used in all MD simulations: Velocities of the system were calculated using the Langevin middle integrator (Zhang *et al.*, 2019). Non-bonded interactions were modelled using the Particle-Mesh Ewald method with a non-bonded cutoff of 10 Å (Darden *et al.*, 1993). Constraints were applied to all bonds between heavy atoms and hydrogen atoms using the SHAKE algorithm (Kr�utler *et al.*, 2001). Bonds and angles of water molecules were restrained using the SETTLE algorithm (Miyamoto and Kollman, 1992). For NpT simulations, the Monte-Carlo barostat was used to enforce a constant pressure of 1 bar. We treated all protons as having a mass of 4.03036 amu, the extra mass being subtracted from adjacent heavy atoms (Feenstra *et al.*, 1999). This, combined with hydrogen bond constraints allowed all simulations to be performed using a timestep of 4 fs, which roughly doubled our simulation speed.
